# Detection of the invasive tomato red spider mite, *Tetranychus evansi* Baker & Pritchard (Acari: Prostigmata: Tetranychidae) in Australia based on morphological identification and DNA sequence analysis

**DOI:** 10.1101/2025.06.11.657326

**Authors:** Danuta Knihinicki, David Gopurenko, Peter S Gillespie, Bernie Millynn, Bernie Dominiak, Louise Rossiter

## Abstract

Australia is an island nation and is free from many of the pests and disease found in other parts of the world. Therefore, Australia developed a national response mechanism to respond to detections of exotic pests. This mechanism includes early detection surveillance and identification services. Tomato red spider mite, *Tetranychus evansi* Baker & Pritchard is a serious pest of solanaceous plants (including tomato, potato and eggplant) in many parts of the world. *T. evansi* was first detected in Australia in August 2013 at Sydney, New South Wales. As part of a response to an exotic incursion, a delimiting survey was initiated. Other *Tetranychus* species exist in Australia and identification techniques were required to identify different species. Submitted survey samples were identified to species level using morphological keys and molecular sequence analysis of the mitochondrial cytochrome *c* oxidase I (COI) gene. Surveillance results of Solanaceae host plants at the time determined it was not technically feasible to eradicate this emergency plant pest (EPP). Additional to blackberry nightshade (*Solanum nigrum* L.), tomato, potato and sweetcorn, *T. evansi* was found damaging native kangaroo apple, *Solanum aviculare* G. Forst., (a new host plant record) and naturalised capeweed, *Arctotheca calendula* (L.) Levyns (Asteraceae). COI sequence analyses confirmed the morphological identifications of three different *Tetranychus* species (including *T. lambi* Pritchard & Baker and *T. ludeni* Zacher) among samples and further identified all twelve *T. evansi* specimens to a single mitochondrial haplotype (clade1; H4) previously reported from climatically diverse locations in Africa, East Asia and the Mediterranean region. Currently, *T. evansi* is considered to be an established plant pest in NSW, so far extensively recorded in the Sydney Basin. Based on the lack of COI sequence variation diversity among the Australian specimens of *T. evansi,* we suggest that this species was likely introduced from a single genetically depauperate source population. This genetic lineage of *T. evansi* established now in Australia was previously reported as invasive in semi-arid and cold regions outside of the species preferred distribution, and laboratory trials have indicated its propensity for increased tolerance of cold climate habitats. This has implications concerning potential for further spread of this lineage of *T. evansi* to a broader range of habitats and plant hosts in Australia.

## INTRODUCTION

Australia is an island nation and is free from many pests that affect their trading partners (Anderson *et al*. 2017; Plant Health Australia 2021). However, the threat of incursion from exotic invertebrate pests and diseases is increasing annually: many incursions occur near major ports of entry because of the volume of freight (100 million tonnes arriving by sea) and the movement of humans in aircraft (9.3 million passengers) (Plant Health Australia 2021). Many exotic pests have established in Australia and adversely impacted many industries and the environment (Gillespie *et al*. 2003; Carnegie & Cooper 2011).

Tomato red spider mite, *Tetranychus evansi* (Baker & Pritchard) is a member of the phytophagous family Tetranychidae which comprises more than 1270 valid species with many being important pests of agriculture. Two-spotted mite, *Tetranychus urticae* Koch, with its global distribution and broad host range, is probably the most well documented (Migeon & Dorkeld 2006–2013), although other invasive species such as *T. evansi* are increasingly becoming the focus because of the wide economic damage caused in new environments. As with other small, plant-feeding mites, often emergent populations of *T. evansi* are only detected when symptoms of visual damage become apparent (Boubou *et al*. 2012; Migeon 2013). Often, continuous feeding by *T. evansi* and production of profuse webbing leads to plant death (Migeon 2013). During the last 20 years with an increase in international trade, *T. evansi* has established itself as an invasive species in new geographic areas (Boubou *et al*. 2011; Navajas *et al*. 2013), a movement that is also reflective of other adventive species groups such as tiny eriophyoid mites (Navia *et al*. 2010).

Originally, *T. evansi* was described from invasive specimens collected on the island of Mauritius in 1960 (Jeppson *et al*. 1975). This mite was first reported from north-eastern Brazil as early as 1915 and deemed to be native to South America (Migeon 2013; Navajas *et al*. 2013), but it is not currently considered to be a pest there (Migeon 2013). In contrast, farmers on the African continent have become increasingly concerned about *T. evansi* becoming a major invasive and widespread pest of tomato crops, especially in Kenya (Toroitich *et al*. 2014) where the species was originally identified in error as *T. urticae* (Knapp *et al*. 2003). Additionally, *T. evansi* is a major pest in the Mediterranean region and recent DNA sequence analyses indicated it was likely to have been introduced there through multiple sources (Boubou *et al*. 2011, 2012; Ferragut *et al*. 2013). This species can easily survive and multiply on many suitable hosts within a wide range of temperatures (Bonato 1999).

Polyphagous *T. evansi* has a problematic taxonomic history and has sometimes been misidentified, especially in the field (Knapp *et al*. 2003). In Mauritius, Moutia (1958) initially considered this species to be *Tetranychus marianae* McGregor. Concerningly, Ferragut *et al*. (2013) reported that after invading new areas, *T. evansi* can eventually displace native or endemic *Tetranychus* species which occur on important solanaceous crops such as tomato, potato, capsicum, eggplant and peanut (Fabaceae). *Tetranychus evansi* can easily be mistaken for other common spider mite species including *T. urticae*, *Tetranychus lambi* Pritchard and Baker, *Tetranychus ludeni* Zacher and *Tetranychus kanzawai* Kishida. This is because one of the commonly used morphological characters for identifying females of *T. evansi* is the overlap in the proximal tactile setae, which is variable in this species. Hence, the standard diagnostic protocol for the European and Mediterranean Plant Protection Organisation by Migeon (2013) appears to be problematic for this character.

Molecular diagnostic tools are available which permit *T. evansi* to be clearly identified and distinguished from other *Tetranychus* species (Knapp *et al*. 2003; Matsuda *et al*. 2013). Furthermore, non-destructive methods of DNA extraction for mites have long been reported (Dabert *et al*. 2008), allowing investigators to perform genetic analyses while also maintaining specimen exoskeletons for downstream morphological analyses and voucher preparations (Skoracka *et al*. 2012). Despite availability of these resources, up to 30% of *Tetranychus* species sequence accessions available at GenBank are taxonomically misidentified (see review by de Mendonça *et al*. 2011). Although problematic for investigation of Tetranychidae systematics in general, none of the species misidentifications indicated by de Mendonça *et al*. (2011) involved overlap with *T. evansi*. This was evidenced at both mitochondrial and nuclear genes as a genetically monophyletic species distinguishable from all other Tetranychidae (Matsuda *et al*. 2013). Furthermore, recent population genetic analyses indicated the presence of two interbreeding genetic lineages of *T. evansi* that minimally differ by 2.37 % and 0.19 % at the mitochondrial cytochrome *c* oxidase I (COI) and nuclear ribosomal intergenic spacer (ITS) loci respectively (Gotoh *et al*. 2009; Boubou *et al*. 2011, 2012). The two lineages (clade I & II as defined by Boubou *et al*. 2011) are native to South America, and both were introduced elsewhere. Multiple introductions originating from different sources in South America have given rise to partially overlapping invasive populations of the two clades in the Mediterranean basin. Conversely, invasive populations in Africa and East Asia have emerged from clade I introductions likely sourced from south-west Brazil (Boubou *et al*. 2011, 2012). Laboratory evidence of increased cold tolerance of the widespread clade I lineage is suggestive of its broader adaptability and may explain its greater invasive range (Migeon *et al*. 2015). Regardless, novel emergences of the species can be examined for their genetic affiliation to the two clades, and this information may be useful for future reference if the two clades differ in their responses to bio-controls or other mitigation practices.

In Australia, *T. evansi* was initially detected on blackberry nightshade, *Solanum nigrum* L., at Port Botany, NSW by Department of Agriculture and Water Resources (DAWR) Central East Region Targeted Surveillance (CERTS) in conjunction with DAWR Operational Science Support (OSS) in August 2013 and identified by DAWR to species level. As part of Australia’s response to the detection of an exotic plant pest, a delimiting survey of potential host plants concentrating on the major host family Solanaceae was immediately conducted.

Here, we report on the statewide survey and the identification techniques using traditional morphological identification and non-destructive DNA sequencing for comparison. By the end of the survey in February 2014, *T. evansi* has only been reported within the Sydney Basin, NSW, where it is considered to be a newly established pest (NSW Department of Primary Industries 2013). Apart from collections on blackberry nightshade, tomato, potato and sweetcorn, *T. evansi* was found causing extensive damage to kangaroo apple, *Solanum aviculare* G. Forst, a flowering, soft-wooded native shrub which grows on the east coast of Australia and New Zealand.

## MATERIAL AND METHODS

### Sampling and preparations

A survey at 238 sites from 92 different localities in NSW was undertaken between August 2013 and February 2014 to determine the presence or absence of *T. evansi* (Fig. 1). The sampling sites were mainly chosen opportunistically but concentrating on those with potential host plants, especially in the genus *Solanum*. Within the Sydney Basin area, the chosen sampling sites radiated outwards from the initial detection site at Botany Bay with an emphasis on the northern, southern and western regions. Other opportunistic sampling, in conjunction with other activities, occurred in other parts on NSW. Plants displaying typical damage symptoms were targeted and, in some instances, mites were observed on leaves in the field (with the aid of a hand lens). Plant samples were collected, placed inside plastic bags and kept cool for transportation back to the laboratory. Prior to detailed examination and with respect to best-practice quarantine protocols, plant samples were placed in the freezer at −4 °C for 24 hours until all potential mites were dead.

**Fig. 1.**
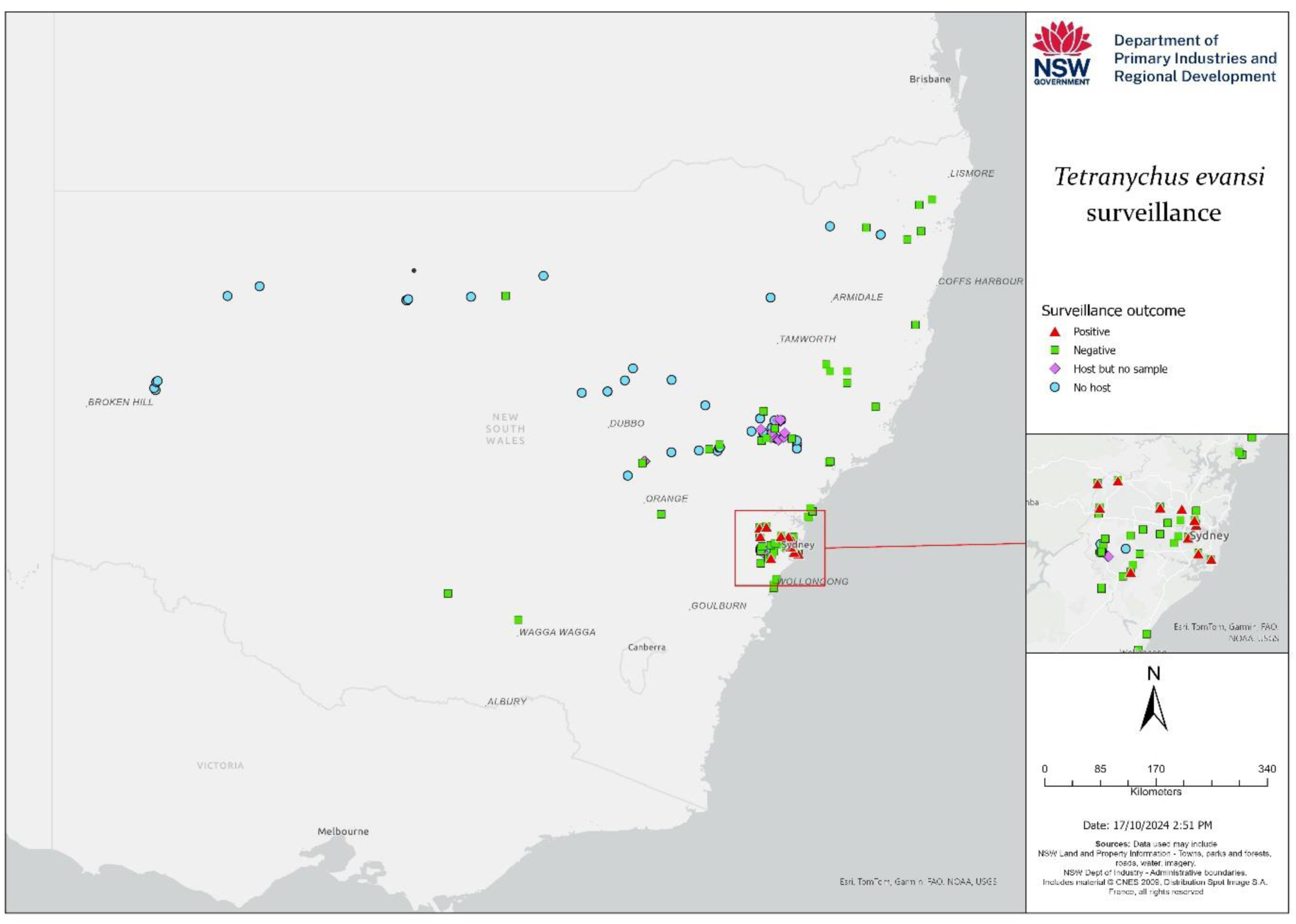
Survey results (August 2013–January 2014) showing presence and absence of tomato red spider mite, *Tetranychus evansi*, and all the sampling sites in NSW. Detailed insert showing presence and absence of *T. evansi* in the Sydney Basin.

Later, mites were collected directly from the plant material by dipping and shaking the foliage in 70% ethanol followed by direct examination under a stereo-microscope. When tetranychids were present in the sample, representatives were slide mounted (see next) and identified as far as possible using morphological techniques. Since traditional morphological identification to species level is based on appearance of the aedeagus, only samples containing males could reliably be determined that way. Hence, this was not possible because sometimes many samples contained only female or immature stages. Molecular diagnosis was employed to determine species identity and as a means of checking morphological identifications (see later).

Slide mounted voucher specimens of *T. evansi* were deposited in the reference collections of the Agricultural Scientific Collections Trust (ASCT), NSW Department of Primary Industries, Orange Agricultural Institute, Orange NSW and Department of Agriculture and Water Resources, Sydney.

### Morphological observations

Individual male specimens were mounted in lateral position on microscope slides directly into Hoyer’s medium (Krantz 1978, p.88; Henderson 2001) enabling a side-view of the aedeagus (required for species level identification in spider mites). Representative females were mounted in the usual dorsal-ventral position. Then, slide mounts were cured at 60 °C in a drying oven for a minimum of 24 hrs, enabling specimens to become translucent. In some instances, specimens were first cleared in Nesbitt’s solution prior to slide mounting without further heating in an oven (J. Otto pers. comm., 19 May 2016). Later, specimens were examined under 40× and 100× (oil immersion) objectives of either an Olympus BX50 compound microscope equipped with phase contrast or a Leica DM2500 compound microscope fitted with differential interference contrast. Species level identification was achieved with the aid of dichotomous keys as provided by Gutierrez and Schicha (1983), Seeman and Beard (2011) and the standard diagnostic protocol published by the European and Mediterranean Plant Protection Organization (Migeon 2013), noting that the overlapping tactile setae on the female legs is a variable character. In the case of samples with female specimens only, the key to major *Tetranychus* groups based on females by Flechtmann and Knihinicki (2002) was used. Migeon and Dorkeld (2006–2013) was consulted for information about the taxonomic nomenclature of *T. evansi* and known host plant species. Images of important morphological characters such as the male aedeagus and variation in female tactile leg setae were captured under 1000× magnification (oil immersion) using a Nikon DS-Fi1 camera.

### DNA sequence analyses

Specimens used for molecular DNA analysis (Table 1) were each allocated a specimen record number for process tracking. DNA from specimens was released by non-destructive digestion in 200 μl of lysis buffer containing 1% Proteinase K at 35 °C for 4 hours (modified from Knölke *et al*. (2005). Following digestion, intact specimens were removed and stored in 70% ethanol and later mounted onto microscope slides following methods described earlier. DNA extraction of specimen digests and subsequent polymerase chain reaction (PCR) preparations followed semi-automated procedures that were reported by Gopurenko *et al*. (2013).

**Table 1.**
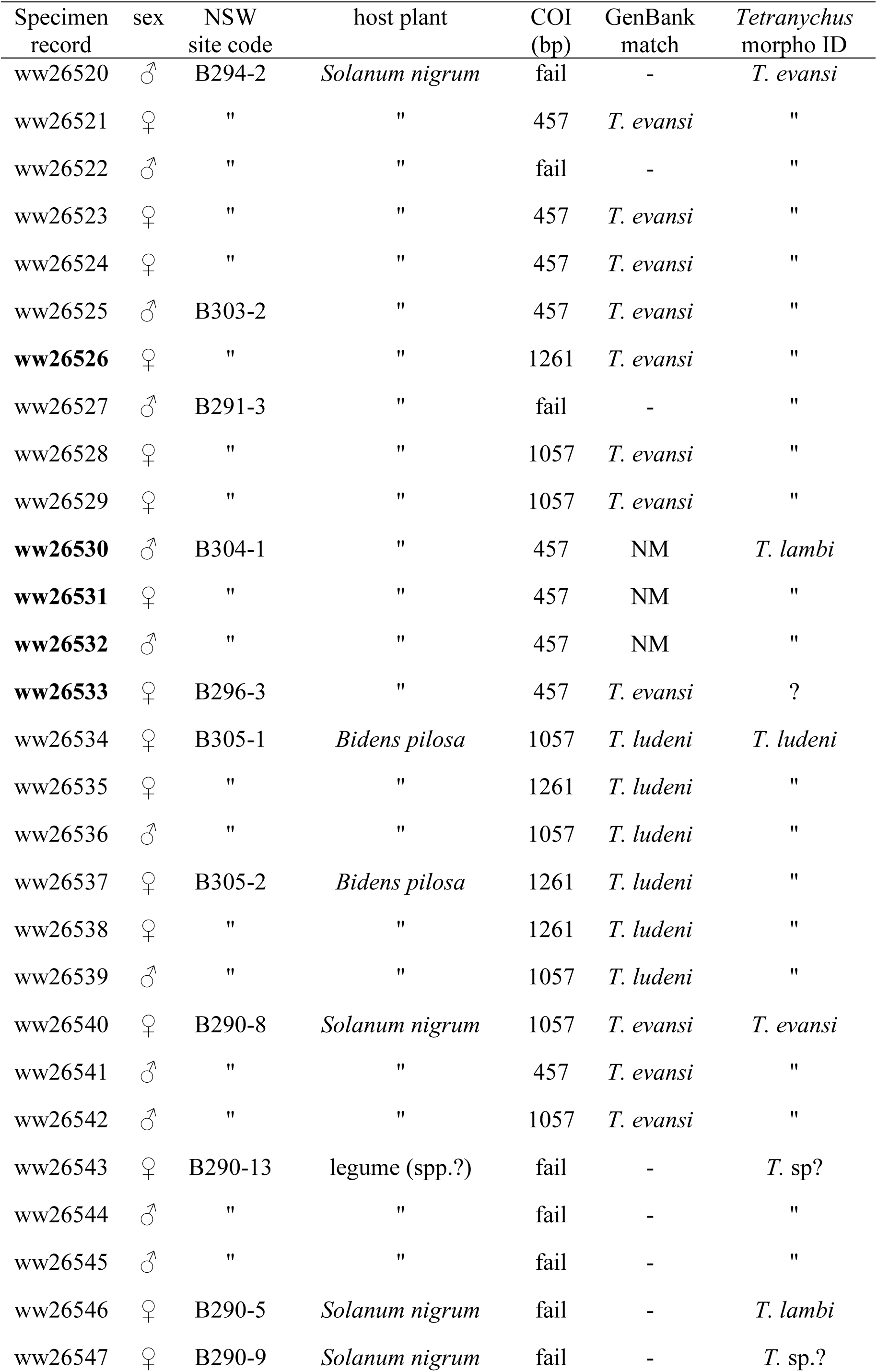
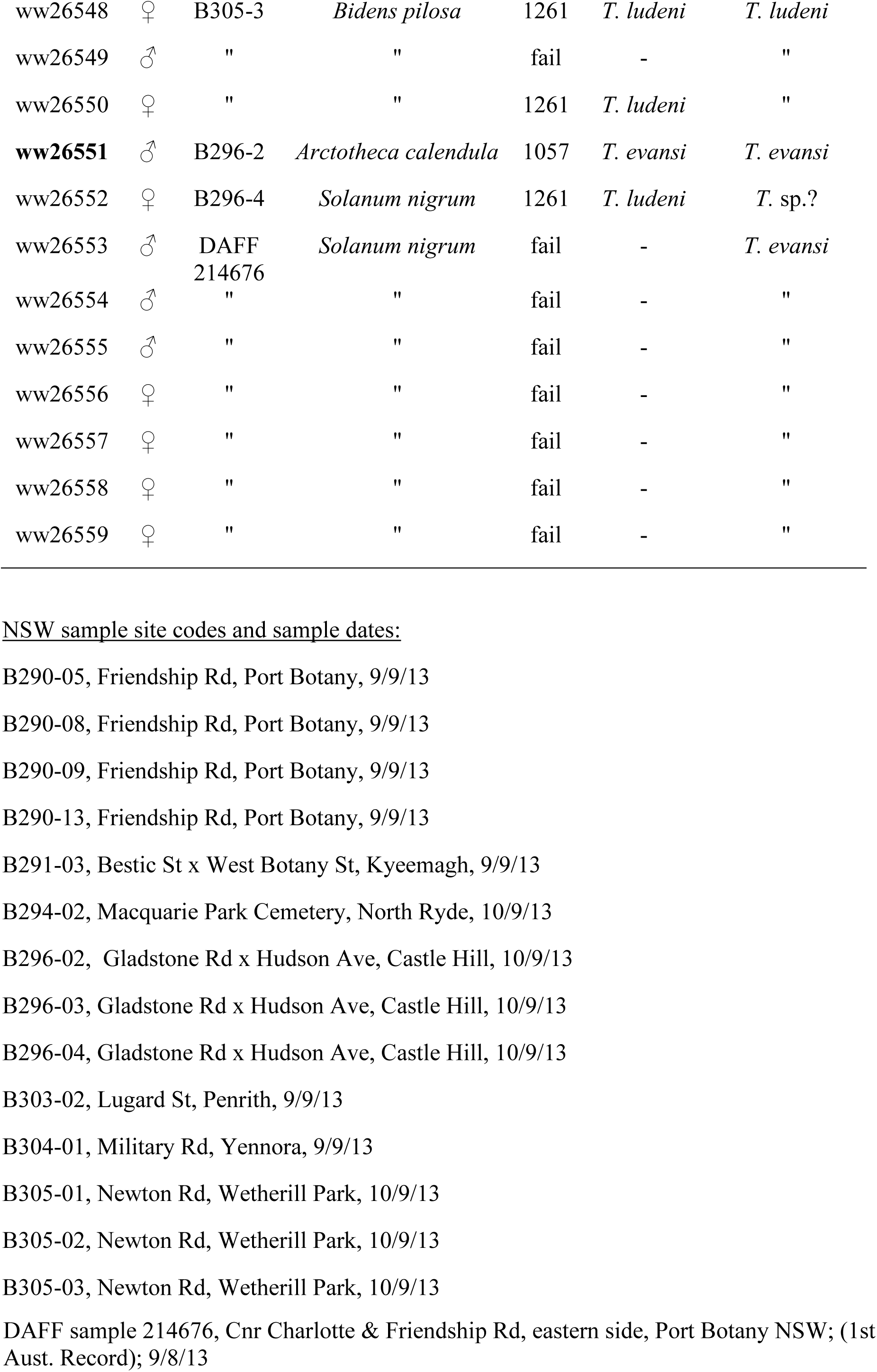
Morphological and genetic identity of *Tetranychus* species sampled at NSW survey sites. Specimen records have been publicly released at BOLD. Specimen sex, host plant, and COI 5’ sequence length (bp) as indicated. GenBank matched COI identifications (> 99% sequence similarity) and morphological identifications of specimens to species as indicated. Slide mounted specimens indicated in bold type. Absence of a genetic match (NM) to existing sequence accessions at GenBank as indicated (searched 16 Sept. 2016).

| Specimen record | sex | NSW site code | host plant | COI (bp) | GenBank match | *Tetranychus* morpho ID |
| --- | --- | --- | --- | --- | --- | --- |
| ww26520 | ♂ | B294-2 | *Solanum nigrum* | fail | - | *T. evansi* |
| ww26521 | ♀ | " | " | 457 | *T. evansi* | " |
| ww26522 | ♂ | " | " | fail | - | " |
| ww26523 | ♀ | " | " | 457 | *T. evansi* | " |
| ww26524 | ♀ | " | " | 457 | *T. evansi* | " |
| ww26525 | ♂ | B303-2 | " | 457 | *T. evansi* | " |
| **ww26526** | ♀ | " | " | 1261 | *T. evansi* | " |
| ww26527 | ♂ | B291-3 | " | fail | - | " |
| ww26528 | ♀ | " | " | 1057 | *T. evansi* | " |
| ww26529 | ♀ | " | " | 1057 | *T. evansi* | " |
| **ww26530** | ♂ | B304-1 | " | 457 | NM | *T. lambi* |
| **ww26531** | ♀ | " | " | 457 | NM | " |
| **ww26532** | ♂ | " | " | 457 | NM | " |
| **ww26533** | ♀ | B296-3 | " | 457 | *T. evansi* | ? |
| ww26534 | ♀ | B305-1 | *Bidens pilosa* | 1057 | *T. ludeni* | *T. ludeni* |
| ww26535 | ♀ | " | " | 1261 | *T. ludeni* | " |
| ww26536 | ♂ | " | " | 1057 | *T. ludeni* | " |
| ww26537 | ♀ | B305-2 | *Bidens pilosa* | 1261 | *T. ludeni* | " |
| ww26538 | ♀ | " | " | 1261 | *T. ludeni* | " |
| ww26539 | ♂ | " | " | 1057 | *T. ludeni* | " |
| ww26540 | ♀ | B290-8 | *Solanum nigrum* | 1057 | *T. evansi* | *T. evansi* |
| ww26541 | ♂ | " | " | 457 | *T. evansi* | " |
| ww26542 | ♂ | " | " | 1057 | *T. evansi* | " |
| ww26543 | ♀ | B290-13 | legume (spp.?) | fail | - | *T. sp?* |
| ww26544 | ♂ | " | " | fail | - | " |
| ww26545 | ♂ | " | " | fail | - | " |
| ww26546 | ♀ | B290-5 | *Solanum nigrum* | fail | - | *T. lambi* |
| ww26547 | ♀ | B290-9 | *Solanum nigrum* | fail | - | *T. sp.?* |
| ww26548 | ♀ | B305-3 | *Bidens pilosa* | 1261 | *T. ludeni* | *T. ludeni* |
| ww26549 | ♂ | " | " | fail | - | " |
| ww26550 | ♀ | " | " | 1261 | *T. ludeni* | " |
| **ww26551** | ♂ | B296-2 | *Arctotheca calendula* | 1057 | *T. evansi* | *T. evansi* |
| ww26552 | ♀ | B296-4 | *Solanum nigrum* | 1261 | *T. ludeni* | *T. sp.?* |
| ww26553 | ♂ | DAFF<br>214676 | *Solanum nigrum* | fail | - | *T. evansi* |
| ww26554 | ♂ | " | " | fail | - | " |
| ww26555 | ♂ | " | " | fail | - | " |
| ww26556 | ♀ | " | " | fail | - | " |
| ww26557 | ♀ | " | " | fail | - | " |
| ww26558 | ♀ | " | " | fail | - | " |
| ww26559 | ♀ | " | " | fail | - | " |

NSW sample site codes and sample dates:
B290-05, Friendship Rd, Port Botany, 9/9/13
B290-08, Friendship Rd, Port Botany, 9/9/13
B290-09, Friendship Rd, Port Botany, 9/9/13
B290-13, Friendship Rd, Port Botany, 9/9/13
B291-03, Bestic St x West Botany St, Kyeemagh, 9/9/13
B294-02, Macquarie Park Cemetery, North Ryde, 10/9/13
B296-02, Gladstone Rd x Hudson Ave, Castle Hill, 10/9/13
B296-03, Gladstone Rd x Hudson Ave, Castle Hill, 10/9/13

A 1261 bp portion of the mitochondrial cytochrome *c* oxidase subunit I (COI) gene, encompassing partially overlapping *Tetranychus* spp. COI sequences available at GenBank, was targeted for PCR amplification and sequencing. Independent PCRs using reverse primer bcdR04 (Dabert *et al*. 2008) in conjunction with forward primers LF1 (Hebert *et al*. 2004) or CI-J-1718 (Simon *et al*. 1994) targeted full length (661 bp) and partial (457 bp) portions of the COI DNA barcode region (Hebert *et al*. 2003) respectively. Primers C1-J-2183 (Simon *et al*. 1994) and Cl-N2776 (Simon *et al*. 2006) targeted a 592 bp COI fragment downstream and adjacent to the DNA barcode region. All primers incorporated 5’ M13 tails to facilitate sequencing. A single thermal profile was used for PCR amplifications: 94° C for 2 min; 40 cycles of 94° C for 30 s, 50° C for 30 s, 72° C for 1 min; 72° C for 5 min; storage at 4° C. PCR products were checked for expected fragment size against E-Gel size marker (Invitrogen), following electrophoresis through a 1.5% agarose gel in 1% TAE buffer. PCR products were sent to the Australian Genome Research Facility (Brisbane) for purification and bidirectional sequencing using an Applied Biosystems 3730*xl* DNA Analyzer. Sequences were quality checked and assembled using Lasergene SeqMan Pro ver. 8.1.0(3) (DNASTAR Inc., Maddison, WI, USA). BioEdit ver. 7.0.9.0 (Hall 1999) was used for primer truncations and final sequence alignment against a reference *T. ludeni* sequence (GenBank accession KJ729018) reported by Chen *et al*. (2014). Sequences were accessioned at GenBank (KX281694 - KX281717) and released at the Barcode of Life Data Systems (BOLD) (Ratnasingham & Hebert 2007) as an online dataset with associated specimen records (dx.doi.org/10.5883/DS-RSMITE). Sequences were queried for species identity against pre-existing online sequence accessions (25/01/2017 & retrospectively 13/01/2025) at GenBank and BOLD. Pair-wise genetic distance relationships among specimen sequences and closest matching accessions at GenBank [including *T. evansi* haplotypes H1-H10, (Boubou *et al*. 2011)] were estimated and assessed as a Neighbour Joining (NJ) distance tree (Saitou & Nei 1987) as implemented in MEGA 6.0 (Tamura *et al*. 2013). The tree was rooted on outgroup *Tetranychus urticea* (accessions AB736077 & AB736080). Missing nucleotide sites among sequences were excluded using a pair-wise deletion option in MEGA. Clades were assessed for significance by bootstrapping (10,000 replicates).

## RESULTS

In summary, the nomenclature was reported by Migeon & Dorkeld (2006–2013). The world-wide distribution was reported by Migeon & Dorkeld (2006–2013) including Africa, Europe, parts of Oceania, and the Americas. Around the world, *T. evansi* was recorded on at least 125 plant species belonging to 33 families (Migeon & Dorkeld 2006–2013). These aspects will not be covered further on our paper.

### *Tetranychus evansi* identifications in New South Wales, Australia

The first Australian records of this species (two males) were sampled 09/October/ 2013 from Port Botany in Botany Bay, Sydney. Of the 92 locations (238 sites) sampled during the 2013/2014 delimiting survey, *Tetranychus evansi* were identified at 20 locations (31 sites) in the Sydney basin (Figure 1) and not elsewhere. Note that sampled sites included public areas, industry zones and private properties; site sample details may be obtained by contacting the corresponding author.

### Relation to plant hosts

*Tetranychus evansi* was found on the following host plants: *Solanum aviculare* (1 site), *S. lycopersicum* (1), *S. nigrum* (21), *S. sp.* (5), *S. tuberosum* (1), *Zea mays* (2) and *Arctotheca calendula* (1). The pest was present on both sides of leaves in heavy infestations with a preference for undersides near the veins (Fig. 2). Leaves soon become yellowish-white, taking on a mottled, silver-like appearance. Extensive webbing is produced when populations reach high levels and plants often look mummified. In severe attacks, leaf drop occurs followed by rapid plant death (NSW Department of Primary Industries 2013).

**Fig. 2.**
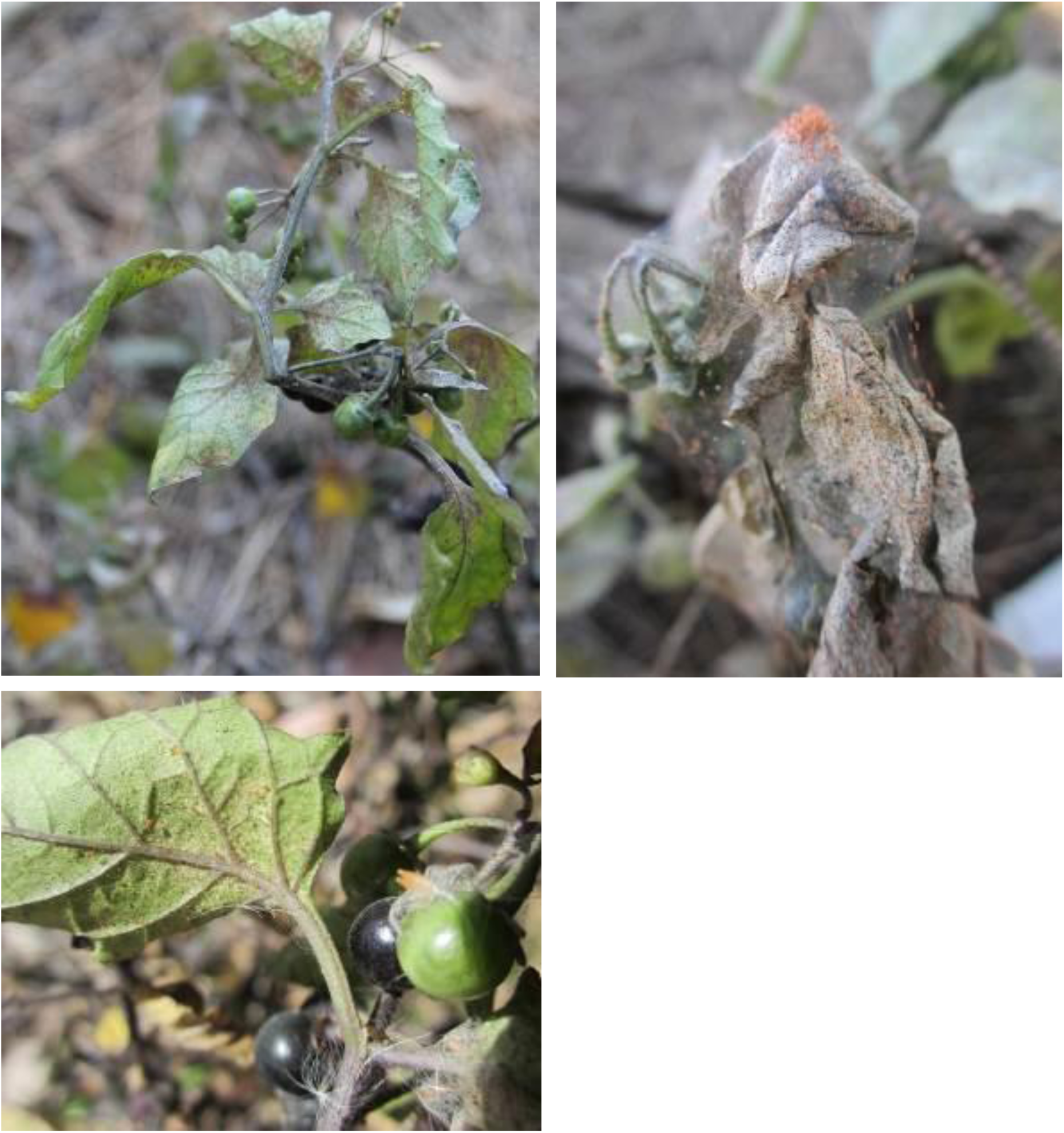
Images of tomato red spider mite, *Tetranychus evansi*, damaging blackberry nightshade, *Solanum nigrum*, in Sydney, NSW.

### Diagnosis

*Tetranychus evansi* belongs to the *desertorum* species group as characterised by Pritchard and Baker (1955): all four proximal setae are in line with the proximal pair of duplex setae on tarsus I; female with dorsomedian empodial spur being tiny or absent. The shape of the male aedeagus is the most important single character for distinguishing *T. evansi* morphologically. As described by Seeman and Beard (2011), the aedeagus is knobbed with a pointed anterior projection and a posterior projection which has a blunt hook. Depending on the focus and angle, the shape of this hook-like structure appears to change.

### Remarks

The records of *T. evansi* listed here are the first to be officially documented from Australia. According to the most recent review by Seeman and Beard (2011), *T. ludeni* a species which is common in NSW, is the only other species from the *desertorum* group which is so far known to occur in Australia.

Native kangaroo apple, *S. aviculare*, is a new host plant record for *T. evansi*. This plant appears to be severely affected (Kearney & Kearney, 2014). In addition, according to Migeon & Dorkeld (2006–2013), the record from Australia is the first for *T. evansi* on capeweed, *Arctotheca calendula* (L.) Levyns (Asteraceae): a single male specimen was confirmed using both morphological and molecular identification.

### Other species identified

Apart from *T. evansi*, four other spider mite species were commonly identified on the various plant species examined during the survey, namely *T. lambi*, *T. ludeni, T. urticae* and *T. kanzawai*. Also, other mites in the families Eriophyidae, Phytoseiidae and Tenuipalpidae were found.

Of interest is that Dr Ray Kearney (pers. comm. January 2014) as part of his observations on *T. evansi* infesting *S. aviculare* at Lane Cove, NSW, found natural enemies such as ladybird beetle larvae (Coleoptera: Coccinellidae) associated with this mite (Kearney & Kearney, 2014). Overseas, Fiaboe *et al*. (2007) and Britto *et al*. (2009) documented the bionomics, predation and reproductive output of the naturally occurring ladybird beetle *Stethorus tridens* Gordon preying on *T. evansi* on solanaceous plants both in the laboratory and field to assess potential for controlling *T. evansi* in tomato crops. The Stethorini have previously been documented as important specialist predators of spider mites (Biddinger *et al*. 2009).

### DNA sequence analyses

Partial COI sequences ranging from 457–1261 bp were obtained from 24 sampled NSW mites (Table 1). Translated sequences were free of indels, stop codons and other symptoms of pseudogene presence. Three haplotypes were identified among the sequences, and these differed by 11.09–2.90 % uncorrected difference. BLAST queries identified one haplotype (present in 12 mites) with 99–100 % similarity to existing *T. evansi* COI sequence accessions. A second haplotype (in 9 mites) genetically matched (> 99 %) to *T. ludeni.* The remaining haplotype was found in three mites that were identified by morpho-analyses as *T. lambi*. This 3^rd^ haplotype was unmatched at GenBank or BOLD sequence repositories and could not be genetically identified to species, though its closest similarity at both repositories (< 91 %) was exclusively to *Tetranychus* species. No pre-existing *T. lambi* COI sequences were available in GenBank or BOLD for comparison to the haplotype (searched 16 Sept. 2016), and therefore the novel haplotype accession reported here represents a first COI sequence entry to GenBank and BOLD for this species. Neighbour Joining distance analysis (Fig. 3) of the NSW mite sequences and comparable closest matching GenBank sequence accessions indicated NSW specimens were grouped in three strongly supported (> 97 % bootstrap) monophyletic species clades. Genetic and morphology-based species identifications agreed in all instances, including at 16 females that were initially identified to species by morphology (Fig 3; Table 1). NJ analysis also identified the single *T. evansi* haplotype present in NSW mites as identical to the “clade I, haplotype 4” lineage of *T. evansi* as defined by Boubou *et al*. (2011, 2012) and previously reported from Africa, east Asia and the Mediterranean region.

**Fig. 3.**
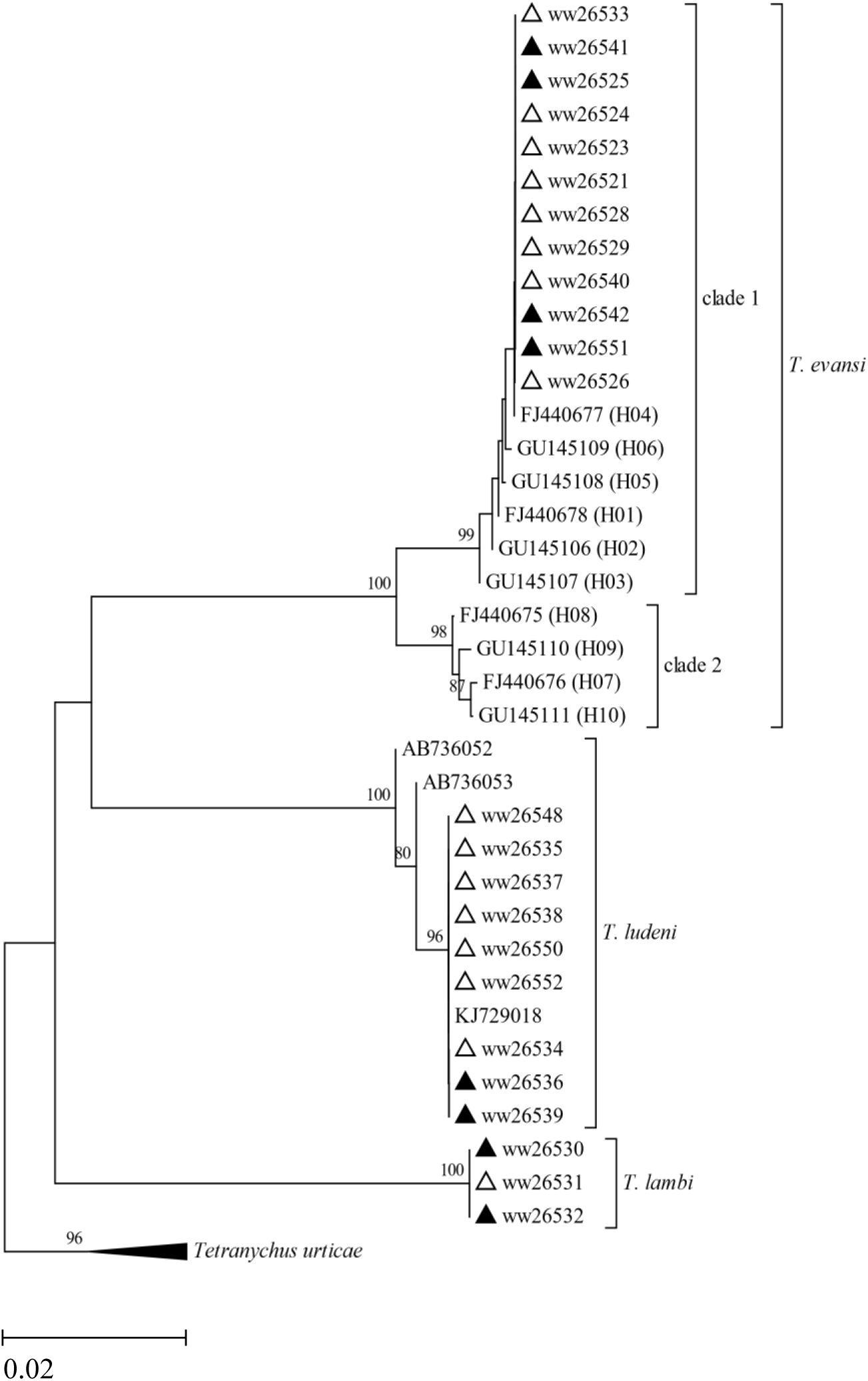
Neighbour joining tree of pairwise COI 5’ genetic distances between 24 sampled NSW *Tetranychus* mites (triangles open = female, shaded = male) and closest matched GenBank sequence accessions. NSW mites labelled with triangle and ww specimen record number (refer Table 1); all other mites labelled with GenBank sequence accession number. *T. evansi* clades I & II and ten haplotypes (H01 -H10) as defined by Boubou *et al*. (2011). Tree rooted at outgroup *Tetranychus urticea* (accessions AB736077 & AB736080). Scale bar equals 2 % COI sequence unweighted distance. Clade bootstrap supports > 70 % as indicated.

## DISCUSSION

Here, we confirmed the presence of an invasive pest mite, *T. evansi*, in Australia, using morphological and molecular analyses of specimens. Based on the presence of the mite among widespread locations in the Sydney basin of NSW, and found on a variety of Solanaceae hosts, we suggest the species has established populations in the region.

In a modelling study by Migeon *et al*. (2009) looking at potential distribution, they found that *T*. *evansi* was most likely a native species of South America since it did not cause major economic damage there. Its presence in Eurasia and North America may be less widespread because of colder climatic conditions. However, in warmer geographic zones, *T. evansi* can reproduce very quickly giving rise to large outbreak populations which can cause significant damage (Soto *et al*. 2010). In Africa, yield losses in tomato crops resulting from infestation were estimated to be as high as 90% (Azandémè-Hounmalon *et al*. 2014).

The discovery of *T. evansi* on *S. aviculare* in Australia may be of some concern since a detailed study on the ecological status of this plant species in New Zealand by Weavers (2010) found *S. aviculare* to be in decline. Interestingly, Murungi *et al*. (2011) observed that severity of infestation by *T. evansi*, a major economic pest species in Africa, greatly varied on different species of nightshade (*Solanum* spp.) and was dependent on the hairiness of leaves. Since leaves of *S. aviculare* are soft and glabrous (Weavers 2010), it may be more prone to attack by *T. evansi*. As the mite increases its spread (Fig. 1), heavy infestations are easily recognised by the vast visible webbing covering plants (Qureshi *et al*. 1969).

*Tetranychus evansi* sampled here were evidenced by the COI sequencing as identical to haplotype 4 (clade 1) *T. evansi* as previously described by Boubou *et al*. (2011 & 2012). Based on the presence of a single mitochondrial lineage among sampled specimens, we suggest a single source entry of this species into Australia, although exact origins of that source cannot be pinpointed here. The detection in Sydney is consistent with its status as the largest Australian port for air and sea movements. This haplotype has the broadest global distribution of all invasive *T. evansi* and is present in northern Africa and separate regions of eastern Asia (Japan, Taiwan) and the Mediterranean basin. The haplotype was not previously identified to any location in South America, perhaps reflecting paucity of earlier native distribution sampling for this species. Regardless, due to its derived genealogical position relative to the inferred ancestral clade 1 haplotype (H01) present in the south-west of Brazil (refer Fig. 2 in Boubou *et al*. 2011), we suspect that it shares a similar native distribution to that and other clade 1 haplotypes. Accordingly, potential biocontrol agents of the species from that region of South America should be considered for future mitigation of the species in Australia. Alarmingly, haplotype 04 was earlier noted (Boubou *et al*. 2011) as invasively present in global habitats beyond the modelled climatic borders for the species. Specifically, the haplotype was present at the margins of semi-arid habitats in Africa, and other regions (within Japan and Taiwan) predicted to be stressful to the species due to extremes of cold and humidity. Laboratory evidence of increased cold tolerance in mites belonging to the widespread clade I lineage is suggestive of their broader adaptability (Migeon *et al*. 2015). Clearly, this has implications concerning the potential for further population expansion by this invasive species into interior Australia because of its preferred environmental habitats and the semi-arid and colder higher-latitude regions of Australia, as earlier predicted by Meynard et al. (2013). This concern is amplified, by our findings that the species is using novel Solanaceae and Asteraceae host plants; entry of *T. evansi* further into diverse Australian habitats may allow it to expand its host plant usage.

## ACKNOWLEDGEMENTS

Many colleagues within NSW Department of Primary Industries and Regional Development and DAWR contributed to this work through the collection and washing of plant samples and are especially thanked (in no particular order): Holger Löcker, Rosy Kerslake, Chris Bloomfield, Michelle Rossetto, Catherine Phillips, Karren Cowan, Michael Priest, Michael Pogson, Cecilia Lawler, Deirdre Gunning, Birgit Löcker, Kathy Gott, Adrian Nicholas, Chris Anderson, Jonathon Lidbetter, Terry Osbourne, Chris McIvor, Darryl Cooper, David Deane, Genevieve Leonard, Sarah Sullivan, Angus Falla, Samantha Hill, Jordan James, Paul Jennings, Daniel Armstrong and Zeljko Begic. Also, we especially thank Ray & Elma Kearney, Sydney, for providing information and specimens from *S. aviculare*. Additionally, we are very grateful to Jennifer Beard (Qld. Mus) for confirming the initial morphological diagnosis of *T. evansi* following its discovery and help from DAWR arachnologist Jürgen Otto. We appreciate reviews of this work graciously provided by NSW DPIRD staff, Lauren Drysdale & Katie Robinson. Funding for molecular work was provided by the NSW *BioFirst initiative* grant to the NSW Agricultural Genomics Centre. David McKenzie, husband of Danuta Knihinicki, encouraged the posthumous finalisation of this manuscript for release and or publication.

